# Inflections in adolescent cortical maturation signal psychopathology risk

**DOI:** 10.64898/2026.10.08.755024

**Authors:** Jacqueline A. Clauss, Dongmei Zhi, Sofia A. Perdomo, Richard Dear, Keiko Kunitoki, Dylan E. Hughes, August T. Blum, Kristina A. Kane, Safia Elyounssi, Bob Cooper, Eline Laurent, Oren Bazer, Louis N. Vinke, Sarah L. O’Dor, Elizabeth A. Flook, Clas Linnman, Michael D. Kritzer-Cheren, Daphne J. Holt, Phil H. Lee, Erin C. Dunn, Hang Lee, Theodore D. Satterthwaite, Alysa E. Doyle, Deanna M. Barch, Jodi M. Gilman, Petra E. Vértes, Edward T. Bullmore, Joshua L. Roffman

## Abstract

Cerebral cortical maturation supports emotional, social, and cognitive development during adolescence^1^. Neurodevelopmental theories of psychiatric illnesses, which peak in onset around age 18^2^, implicate subtle alterations in adolescent cortical development^3^. However, mechanistic links between cortical maturation and emergent psychopathology have remained elusive. This gap likely reflects a need for more nuanced understanding of cortical development^4^, larger, longitudinal samples powered to detect small effects,^5^ and more uniform image quality control^6^. Using 29,482 structural MRI scans from 11,720 youth in the Adolescent Brain Cognitive Development (ABCD) study, we demonstrate reproducible differences in the contour and tempo of maturation across the cortical surface. Regional trajectories are robust to image quality variance and align with increasing functional specialization over time. Thicker cortex in early-maturing sensory regions predicts psychiatric symptoms as early as four years prior to their emergence, a pattern that subsequently spreads to association cortex. Overlap with spatially and temporally organized gene co-expression networks implicates prenatal patterning of this sequence of maturation and its links to psychopathology. These findings present a novel, versatile neurodevelopmental framework for cortical maturation during adolescence that sequentially implicates sensory and association cortex in risk for emergent psychiatric symptoms.

## Introduction

Thinning of the cerebral cortex through childhood and adolescence is a hallmark of brain maturation that underlies emotional, social, and cognitive development. Thought to reflect a combination of synaptic pruning and expansion of white matter tracts^7,8^, cortical thinning can be measured reliably in vivo using structural magnetic resonance imaging (sMRI). Importantly, numerous mental illnesses, including mood and psychotic disorders, begin to emerge during adolescence and early adulthood in tandem with these cortical changes, suggesting their interdependence^9^. Indeed, several serious mental illnesses, most notably schizophrenia, have associated with distributed reductions in cortical thickness that are postulated to reflect altered neurodevelopment^10^.

However, associations between cortical thinning and psychopathology have remained equivocal, reflecting several experimental limitations. First, the basic spatial and temporal patterning of cortical maturation is incompletely understood, even in normative samples – which, for example, have demonstrated linear contours in some studies, and non-linear contours in others^4,11–13^. Variation in image quality, which is particularly sensitive to motion artifacts among young participants, may contribute to these discrepancies^4,6^. Second, brain-behavior associations are often subtle, requiring sample sizes into the thousands to detect reliably, and contributing to inconsistent findings in smaller samples^5^. Third, cortical maturation and emergence of psychopathology are each highly dynamic processes that limit the value of cross-sectional studies. Mental illnesses such as major depressive disorder and schizophrenia frequently arise from nonspecific symptoms in adolescence (such as anxiety or attention problems) that are poorly aligned with clinical diagnostic frameworks^14^. These early symptoms may also exert downstream effects on cortical maturation itself^15^, complicating causal inference.

Here, we overcome these challenges by using repeated-measures, high quality sMRI and dimensional measures of psychopathology, from more than 11,000 youth in the Adolescent Brain and Cognitive Development (ABCD) Study^16^. We first sought to characterize spatial and temporal maturation patterns across the cortex, and to assess their relevance to fundamental developmental processes such as functional specialization and sex differences. On this foundation, we then identified and sequenced relationships between cortical maturation and emergent symptoms, and tested their congruence with developmentally discrete gene expression programs.

## Results

The ABCD Study enrolled 11,880 children for a follow-up period of ten years, including biennial MRI scans and annual parent-reported psychopathology assessed via the Child Behavior Checklist (CBCL; **Fig 1a**). The current analysis used ABCD Data Release 6.0 (N=11,820 with available scans), comprising data from Baseline (ages 9-10) through Year Six (ages 15-16). We excluded scans by timepoint if they had unacceptable image quality (surface hole number, SHN > 62) or missing covariates (totaling 2.6% of available scans and 0.8% of individuals; **Table S1a**). Age-thickness trajectories were first assessed in an optimized longitudinal sample (**Fig. 1b, Table S1b)**, including all 1,476 individuals with a complete, high quality longitudinal dataset (complete covariate data and T1 scans at ages 9-10, 11-12, 13-14, and 15-16, each scan with SHN of < 30, a threshold derived by manual review of >12,000 volumes of sMRI data^6^). All remaining individuals with at least one, and up to four, scans of adequate quality (SHN < 63^6^) were included in the remainder sample (10,244 individuals), to assess reproducibility under less restrictive quality and completeness criteria (**Table S1b**).

**Fig. 1.**
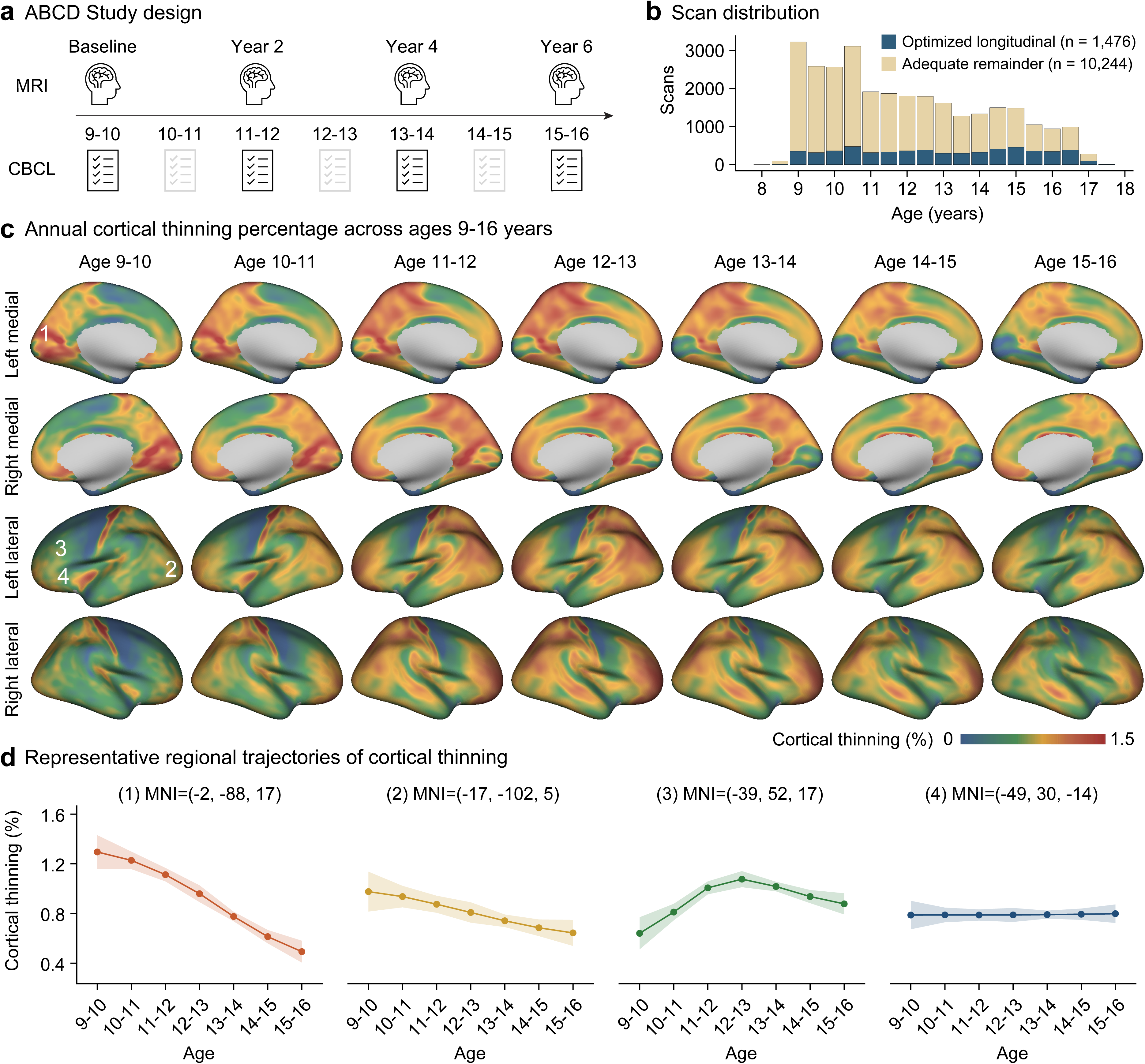
Longitudinal cortical thinning from late childhood to adolescence. **(a)** Design of the Adolescent Brain Cognitive Development (ABCD) Study. The ABCD Study enrolled 11,880 children at ages 9-10, with prospectively acquired biennial structural magnetic resonance imaging (sMRI) scans, and annual parent-reported psychopathology collected from baseline (ages 9–10 years). We included data through the 6-year follow-up (ages 15–16 years). **(b)** Distribution of MRI scans across age. Participants were partitioned into (1) an optimized longitudinal dataset comprising individuals with high-quality scans (SHN < 30) available at all four time points, and complete standard covariate data (*n* = 1,476, see **Methods**), and (2) a non-overlapping adequate remainder dataset including all remaining participants with at least one eligible scan (SHN < 63) and complete covariate data (*n* = 10,244). Bars indicate the number of MRI scans available in each 0.5-year age bin. Scans from the optimized longitudinal and adequate remainder datasets are shown in blue and yellow, respectively. **(c)** Vertex-wise maps of annual cortical thinning across ages 9–16 years, after correction for standard covariates. Maps show annual cortical thinning percentages for each age interval from 9–10 to 15–16 years. Warmer colors denote more cortical thinning; cooler colors denote less thinning. **(d)** Four representative age-related cortical thinning trajectories are shown for regions at Montreal Neurological Institute (MNI) coordinates. See numbers (1-4) in the left-most column in figure (c) for corresponding locations. Clouds indicate 95% confidence intervals (CI).

To provide an overview of age-related thinning trajectories, cross-sectional cortical thickness reductions as a function of integer age (e.g., 9.0 to 9.9 versus 10.0 to 10.9) were mapped across the entire cortical surface in the optimized sample. Thickness values were adjusted for routine biological (sex and age, i.e., to control for age differences within each year-long stratum), technical (intracranial volume, scanner manufacturer), quality control (SHN), and design-related (site, subject ID, family ID) covariates. Marked heterogeneities in both the contour and tempo of age-related thinning trajectories were apparent across the cortex (**Fig. 1c**). Specifically, between ages 9 and 16, distinct cortical regions exhibited (1) early rapid thinning followed by pronounced deceleration, (2) comparatively longer-lasting thinning with later deceleration, (3) an initial acceleration and subsequent deceleration of thinning, or (4) relatively stable thinning rates throughout adolescence (**Fig.1d**).

In parallel, we leveraged repeated-measures sMRI data from the optimized sample to quantitatively characterize age-related cortical thinning trajectories across 68 regions, as parcellated by the Desikan-Killiany atlas^17^. We used a staged approach, contrasting thinning contours based on three (ages 9-14) and then four (ages 9-16) timepoints, to assess stability of contours over time. We used linear mixed-effects models (LMMs) to regress cortical thickness on age, age^2^, and (for six-year models) age^3^, accounting for routine covariates as above (**Methods**). This approach yielded four distinct developmental trajectories across the 68 regions (**Fig. 2**; **Tables S2 and Table S3**; **Fig. S1**) that recapitulated those seen in surface-wide maps (**Fig. 1d**). First, six primary visual regions demonstrated positive quadratic (decelerating) contours through ages 13-14 that persisted through ages 15-16 – consistent with an “early maturing” trajectory. Second, nine regions, primarily in lateral occipital and limbic cortices, transitioned from linear contours through age 13-14 to positive quadratic contours through ages 15-16 – signaling comparatively later maturation (“mid maturing”). Third, 28 regions, predominantly within frontal and parietal association cortices, were best fit by negative quadratic contours though ages 13-14 and cubic (sigmoid) contours through ages 15-16 – reflecting an initial acceleration and later deceleration in cortical thinning (“fast maturing”). Finally, 25 regions, largely spanning transmodal association cortex, demonstrated persistent linear thinning through ages 15-16, with no evidence of acceleration or deceleration (“slow maturing”).

**Fig. 2.**
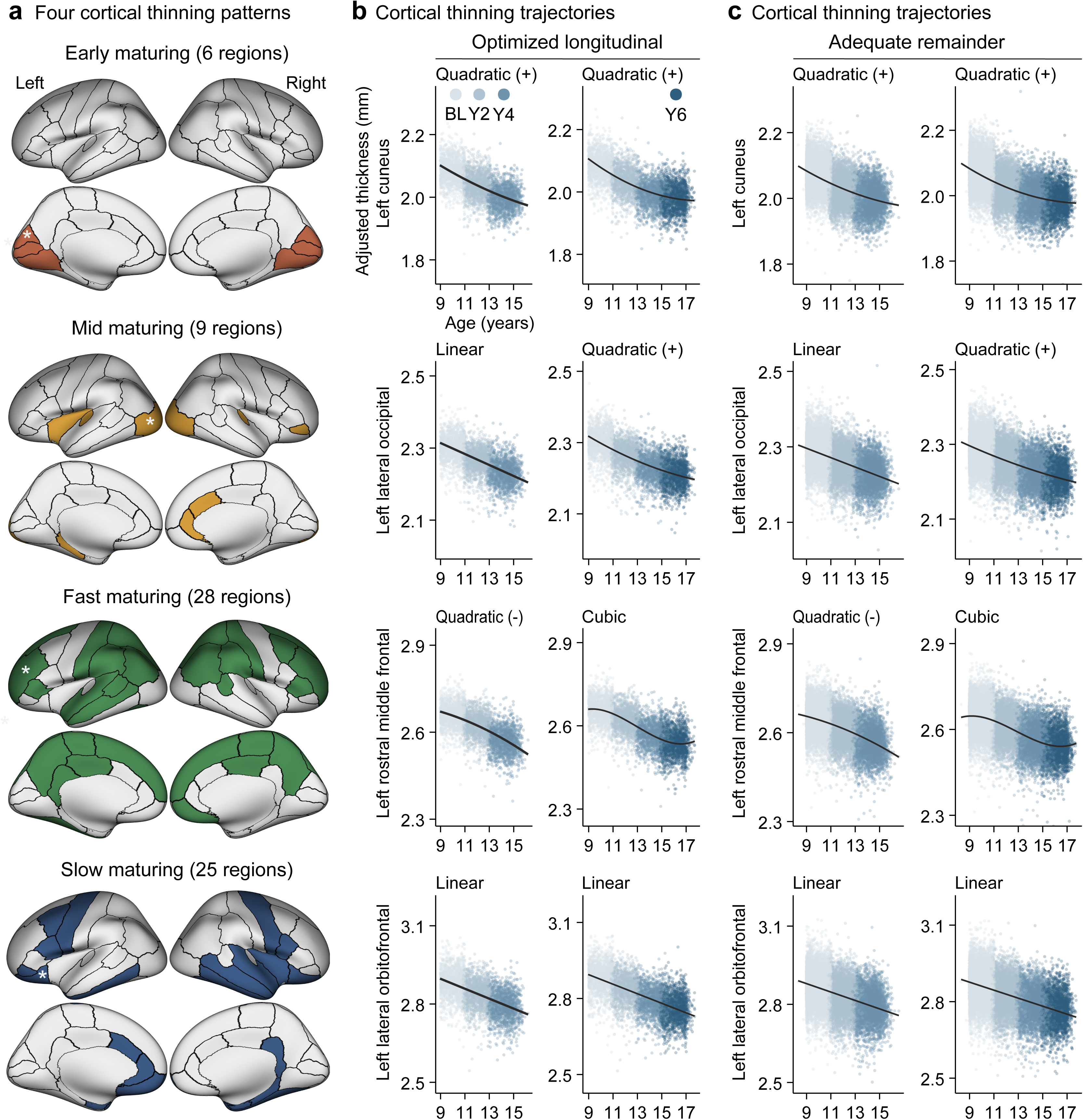
Four cortical thinning patterns. **(a)** Brain maps of the four cortical thinning trajectories: early maturing, mid maturing, fast maturing and slow maturing. Representative regions shown in **(b)** and **(c)** are indicated by white asterisks, which also correspond to the four regions highlighted in Fig. 1 (c) and (d). Early maturing regions were primarily located in the visual cortex; mid maturing regions in the lateral occipital and limbic cortices; fast maturing regions in frontal and parietal association cortices; and slow maturing regions in the temporal, anterior cingulate and precentral cortices. **(b)** Representative cortical thickness trajectories estimated using the first three imaging waves (Baseline, Year 2, Year 4) and all four imaging waves (Baseline through Year 6), in the optimized longitudinal set (*n* = 1,476 individuals, 5,904 scans). Cortical thickness was adjusted for sex, intracranial volume (ICV) and surface hole number (SHN), with participant, family, scanner, and site included as random effects. Each point represents an individual observation color-coded by imaging wave; black lines indicate best-fitting trajectories. Early maturing regions were best fit by positive quadratic trajectories in both analyses. Mid maturing regions transitioned from linear to positive quadratic trajectories following inclusion of the fourth imaging wave. Fast maturing regions transitioned from negative quadratic to cubic trajectories. Slow maturing regions retained linear trajectories in both analyses. **(c)** These patterns largely replicated in the non-overlapping adequate remainder set (*n* = 10,244 individuals, 23,806 scans). In (b) and (c), for visualization only, observations exceeding 5 standard deviations from the regional mean cortical thickness were not shown. The number of observations omitted from visualization ranged from 0–1 and 2–9 across the four representative regions in the optimized and adequate longitudinal datasets, respectively.

These patterns indicate that the specific developmental timing and duration of thickness observations during adolescence are critical to deciphering cortical maturation trajectories. Depending on these parameters, regions may appear to thin linearly (in mid-stages of maturation) or non-linearly (in early and late stages of maturation). However, the observed patterns in every cortical region align with a unifying model of sigmoid-shaped trajectories, of varying onset and duration across regions. Within this six-year window of observation, all three phases (accelerating, linear, and decelerating thinning) were visible only in rapidly maturing cortex (**Extended Data Fig. 1**).

### Generalizability and reproducibility of trajectories

To account for potential influence of socioeconomic and pubertal status, we examined cortical thinning trajectories after additionally adjusting for income-to-needs ratio (INR) and puberty stage in the optimized set. Cortical thinning patterns were largely consistent, with 62 of 68 regions showing the same trajectory types as in the primary analysis (Cohen’s κ □= □0.87, *p*□ <□ 0.001, **Table S4** and **Table S5**). Further, while previous studies in smaller samples^4^ and in those with shorter follow-up^18^ have described accelerated thinning in girls compared to boys in select cortical regions, we observed near-ubiquitous differences. Significant negative linear age-by-sex assigned at birth (SAB) interactions in 66 of 68 regions indicated an accelerated tempo of cortical thinning in girls regardless of contour (**Extended Data Fig. 2**; **Table S6**). This pattern remained highly consistent after additional adjustment for INR and puberty (64 of 68 regions, **Table S7**).

We then further assessed the reproducibility of these trajectories within the adequate remainder dataset (10,244 non-overlapping participants) using the same analytical framework. Across 68 regions, 46 showed the same trajectory type as in the optimized longitudinal dataset (Cohen’s κ□= □0.53, *p*□ = □1.12×10^-12^), with the greatest reproducibility observed for early maturing (6 of 6 regions) and the lowest for mid maturing (2 of 9 regions; **Fig. S1**; **Table S8** and **Table S9**). Among discrepant mid maturing regions, most (five of seven) displayed linear thinning in the remainder sample, suggesting that among these regions, poorer scan quality may obscure the emergence of thickness decelerations. Further, 43 of 68 regions retained consistent trajectories after additional adjustment for INR and puberty in the adequate remainder dataset (Cohen’s κ □= □0.48, *p* □= □1.02×10^-11^, **Table S10** and **Table S11**). Accelerated thinning was again observed in girls, including after additional adjustment for INR and puberty stage (66 of 68 regions, **Tables S12 and S13**).

### Alignment of maturation sequence with functional hierarchy

With most of the cortical surface showing thinning decelerations before age 16, we next sought to more precisely characterize the timing of regional deceleration and compare this sequence to known patterns of cortical hierarchy. We estimated the age of thinning deceleration in each non-linear region by fitting segmented linear models and identifying the breakpoint that minimized the Bayesian information criterion (BIC) in the optimized longitudinal dataset (**Methods**). Breakpoint age varied substantially across the cortex (**Fig. 3a**; **Fig. S3**; **Extended Data Fig. 3**; **Table S14**), ranging from 13.5 to 15.9 years. For example, we observed estimated breakpoints at 14.2 years in the left cuneus, 15.2 years in the left lateral occipital cortex and 15.5 years in the left rostral middle frontal cortex (**Fig. 3b**). Across all regions, cortical thickness continued to decline after the breakpoint, albeit at a reduced rate, indicating a shift from faster to slower cortical thinning.

**Fig. 3.**
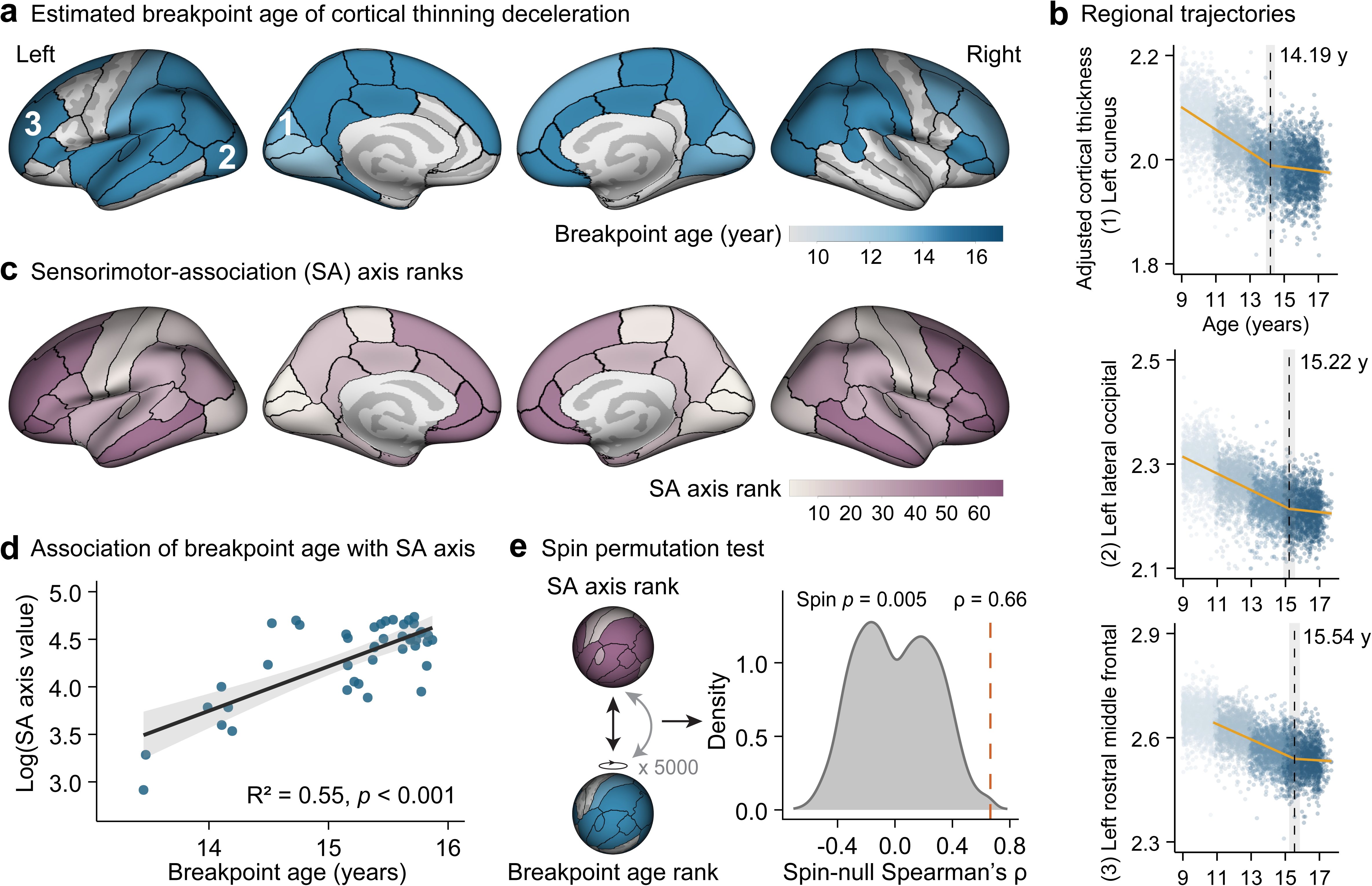
Cortical thinning deceleration follows the sensorimotor-association (SA) axis. **(a)** Brain maps from the optimized sample (*n* = 1,476 individuals, 5,904 scans) showing estimated breakpoint ages of cortical thinning deceleration across regions exhibiting nonlinear developmental trajectories. Numbers denote representative cortical regions shown in Figs. 1 and 2, and Fig. 3 b. **(b)** Breakpoint estimation for representative cortical thickness trajectories, including (1) left cuneus; (2) left lateral occipital cortex; and (3) left rostral middle frontal gyrus. Grey vertical lines indicate estimated breakpoint age with 95% confidence intervals (CIs). Orange lines represent segmented linear model fits before and after the breakpoint. Each point represents an individual observation, color-coded by imaging wave. **(c)** Brain maps showing regional sensorimotor–association (SA) axis ranks. Lower values indicate primary sensorimotor cortex, and higher values indicate transmodal association cortex. **(d)** Association between breakpoint age and the log-transformed continuous SA axis value. Each point represents one cortical region. The solid line indicates the fitted linear regression with 95% CIs (Pearson’s correlation: R^2^ = 0.55, *p* < 0.001). **(e)** Spin testing of the association between breakpoint age rank and the SA axis rank. The grey region represents the null distribution of Spearman’s correlation coefficients generated by spin tests, and the dashed orange line indicates the observed correlation (ρ = 0.66, Spin test *p* = 0.005).

We next examined whether the timing of cortical thinning deceleration was organized along the sensorimotor–association (SA) axis, which measures the region’s relative contributions to lower-order (unimodal sensory and motor) versus higher-order (executive, socioemotional, and mentalizing) functions^1^ (**Fig. 3c**). Breakpoint age was significantly associated with the log-transformed SA axis value (R² = 0.55, *p* < 0.001, **Fig. 3d**), with later regional decelerations occurring progressively towards transmodal association cortex. This spatial organization was confirmed after controlling spatial autocorrelation using spin testing of SA axis rank and breakpoint age rank (*p* = 0.005, 5,000 permutations; **Fig. 3e**).

### Association with dimensional psychopathology

We next examined the relationship between cortical thickness and psychopathology, assessed at ages 9-10, 11-12, and 13-14. To maximize power to detect subtle brain-behavior associations^5,6^, all available cortical thickness data passing minimal quality control (SHN<63) were included in this analysis, and ages 15-16 were not included due to fewer available datasets in the ABCD 6.0 release. Psychopathology was measured using the Child Behavior Checklist (CBCL), which is a parent-reported dimensional measure of symptom severity across depression, anxiety, and somatic complaints (grouped as internalizing symptoms), and rule-breaking and aggression (grouped as externalizing symptoms). As individual symptom domains are highly correlated at ages 9 to 14 (**Extended Data Fig. 4**), we used CBCL total score as the primary clinical measurement, while secondary analyses focused on CBCL internalizing and externalizing symptom scales. We used a hierarchical approach to assess temporal and spatial relationships between cortical thickness and total. First, we tested three competing temporal models of cortex-wide thickness data: 1) “predictive” (earlier cortical thickness predicts later symptoms); 2) “concurrent” (current symptoms associate with current thickness); and 3) “delayed” (earlier symptoms associate with future thickness; **Fig. 4a**). Of these, only the predictive model demonstrated statistical significance, indicating that thickness associated positively with greater, but not concurrent or preceding symptoms (**Table S15a**).

**Fig. 4.**
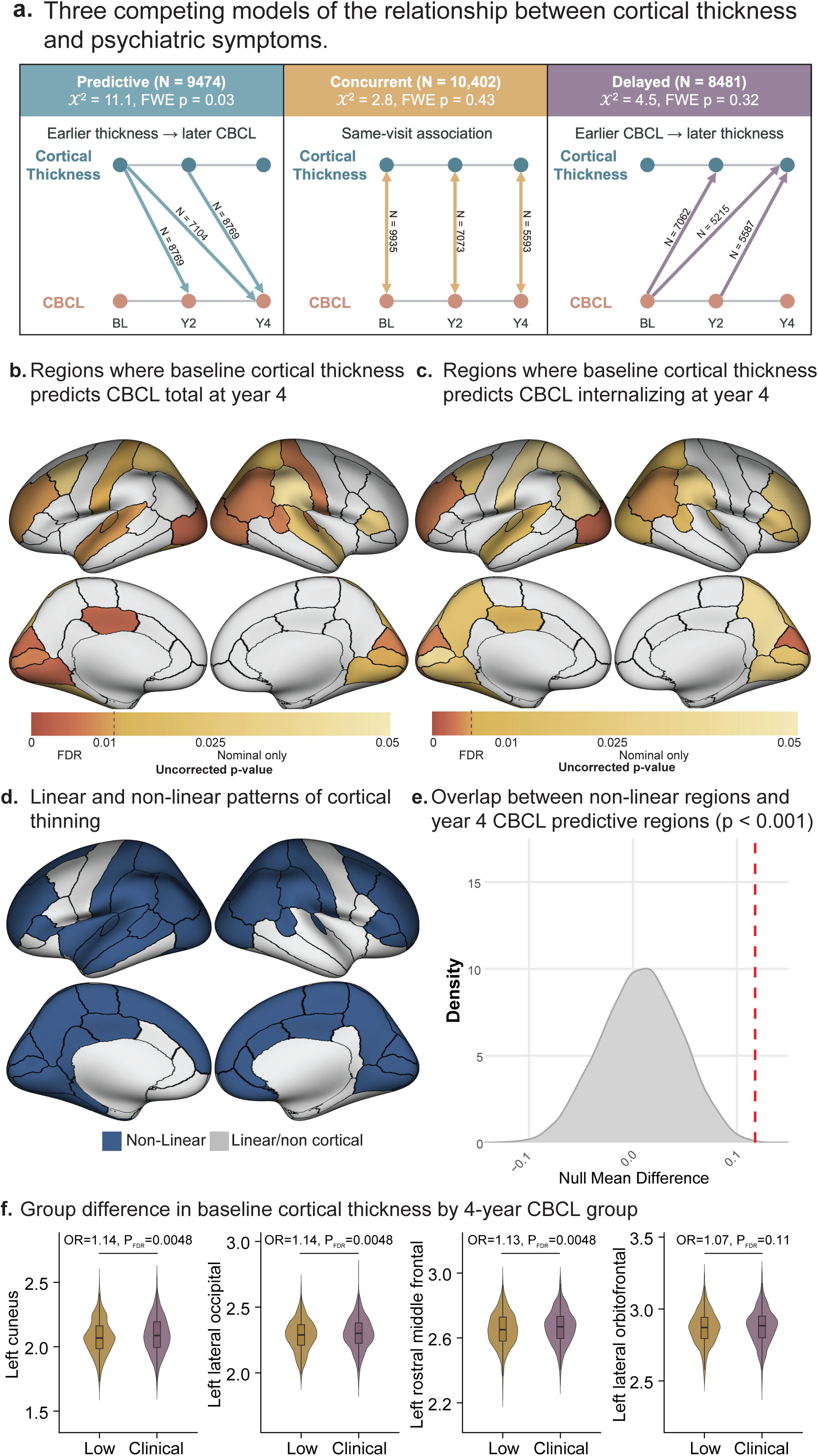
Asynchronous associations between cortical thickness and dimensional symptoms. **(a)** Three models of the relationship between psychopathology and mean cortical thickness were tested: predictive (cortical thinning predicts onset of psychiatric symptoms), concurrent (current cortical thickness predicts current symptoms) and delayed (current symptoms predict later cortical thickness). Only the delayed model was significantly associated with psychopathology (p < 0.05 family-wise error). **(b)** Regions in which cortical thickness at age 9-10 predicted Child Behavior Checklist (CBCL) total score four years later. **(c)** Four-year prediction of CBCL total score largely reflected internalizing symptoms. **(d)** Comparison to (b) suggests that regions with a predictive relationship overlapped with regions showing nonlinear patterns of cortical thinning. **(e)** Spin testing confirmed that regions with nonlinear patterns of thinning were significantly enriched with those where cortical thickness at ages 9-10 predicted CBCL at ages 13-14 (p < 0.001). **(f)** Youth with clinically significant symptoms (*n* = 934) demonstrated thicker cortex than propensity score matched youth from the lowest CBCL tertile (*n* = 934) in regions approaching maturation by age 15-16 (e.g., left cuneus, left lateral occipital, left middle frontal), but not in those with persistent linear thinning (e.g., left lateral orbitofrontal).

We next assessed for variation of this predictive association across 68 cortical regions. Across timepoints, greater cortical thickness in 10 of 68 regions predicted greater subsequent psychopathology (false discovery rate [FDR] corrected *p* < .05); an additional 11 regions were nominally significant (*p* < .05; **Table S16**). These regions included primary visual, somatosensory, and auditory cortices, as well as right posterior parietal and left anterior frontal association cortex. No regions had an inverse association between thickness and subsequent symptoms. Pairwise analyses indicated that these relationships were strongest between cortical thickness at ages 9-10, and psychopathology at ages 13-14 (FDR corrected *p* < .05 in 18 regions, **Fig. 4b; Extended Data Fig. 5; Table S17**). Notably, at age 9-10, the cortical thickness of the left rostral middle gyrus, a fast-maturing region containing higher-order association cortex, did not predict symptoms at age 11-12; but thickness at age 11-12 in this region did predict symptoms at age 13-14 -- consistent with an anterior progression of predictive regions over time as they approach maturation. Overall, predictive relationships were driven primarily by internalizing, rather than externalizing symptoms (**Fig. 4c; Table S15b; Table S18 and Table S19**). Results were robust to image-quality thresholds: effect sizes were highly consistent when analyses were restricted to high-quality scans (SHN <30) compared with analyses including scans passing minimal quality control (SHN <63; R > 0.93; **Extended Data Fig. 6**).

We then examined whether regions in which cortical thickness predicted subsequent psychopathology shared common developmental trajectories. In analyses across timepoints, and in individual timepoint pairwise comparisons, predictive regions occurred only among those with nonlinear trajectories—i.e., early, mid, and fast maturing (**Fig. 4d**); this pattern was confirmed after accounting for spatial autocorrelation using spin testing (**Fig. 4e**). These findings suggest that, in early adolescence, regions that signal future psychopathology are constrained to those that have shown evidence of maturation by age 16; conversely, thickness in regions without evident decelerations (i.e., slow maturing) did not predict psychopathology, at least within the present age window.

Finally, to more directly assess the clinical relevance of these findings and to further reduce risk of confounding bias, we conducted post hoc analyses comparing adolescents with clinically significant symptoms at ages 11-12 or 13-14 (CBCL total ≥60) with participants with CBCL total scores in the lowest third who were propensity-score matched on routine covariates, INR, and pubertal stage (n = 934 per group, Table S20). Of the 18 regions in which thickness at age 9-10 significantly predicted age 13-14y CBCL total score in the full sample, 14 also showed significant differences in age 9-10 thickness between the clinical and matched comparison groups (*p* < .05, FDR; OR = 1.12–1.17; **Fig. 4f**; Table S21).

### Alignment of predictive regions with gene expression profiles

Genetic variation contributes strongly to both cortical thickness and dimensional child psychopathology: single nucleotide polymorphism-based heritability estimates for cortical thickness are as high as 25%^19^, and as high as 18% for CBCL^20^. Further, evidence from postmortem human brain studies demonstrates that gene expression across the cortex is spatially and temporally patterned, with specific gene co-expression networks overlapping with polygenic risk for neurodevelopmental disorders^21,22^. Thus, we next determined whether three previously identified spatially- and temporally-patterned gene co-expression networks align with regions where thickness predicted psychopathology: C1, with peak expression in primary somatosensory cortex and during prenatal life; C2, with peak expression in intermediate sensory-association cortex and during early childhood; and C3, with peak expression in heteromodal association cortex and during adolescence^22^. Using spin testing, we examined spatial overlap between cortical regions that predict later psychopathology (using all time points) and region-specific loading for prenatal, early childhood, or adolescent gene expression patterns. Distribution of predictive regions overlapped significantly with those with higher prenatal (C1), but not early childhood (C2) or adolescent (C3) gene expression (**Fig. 5**). These findings suggest that relationships between cortical maturation and psychopathology in mid-adolescence may reflect prior genetic programming from as early as prenatal life.

**Fig. 5:**
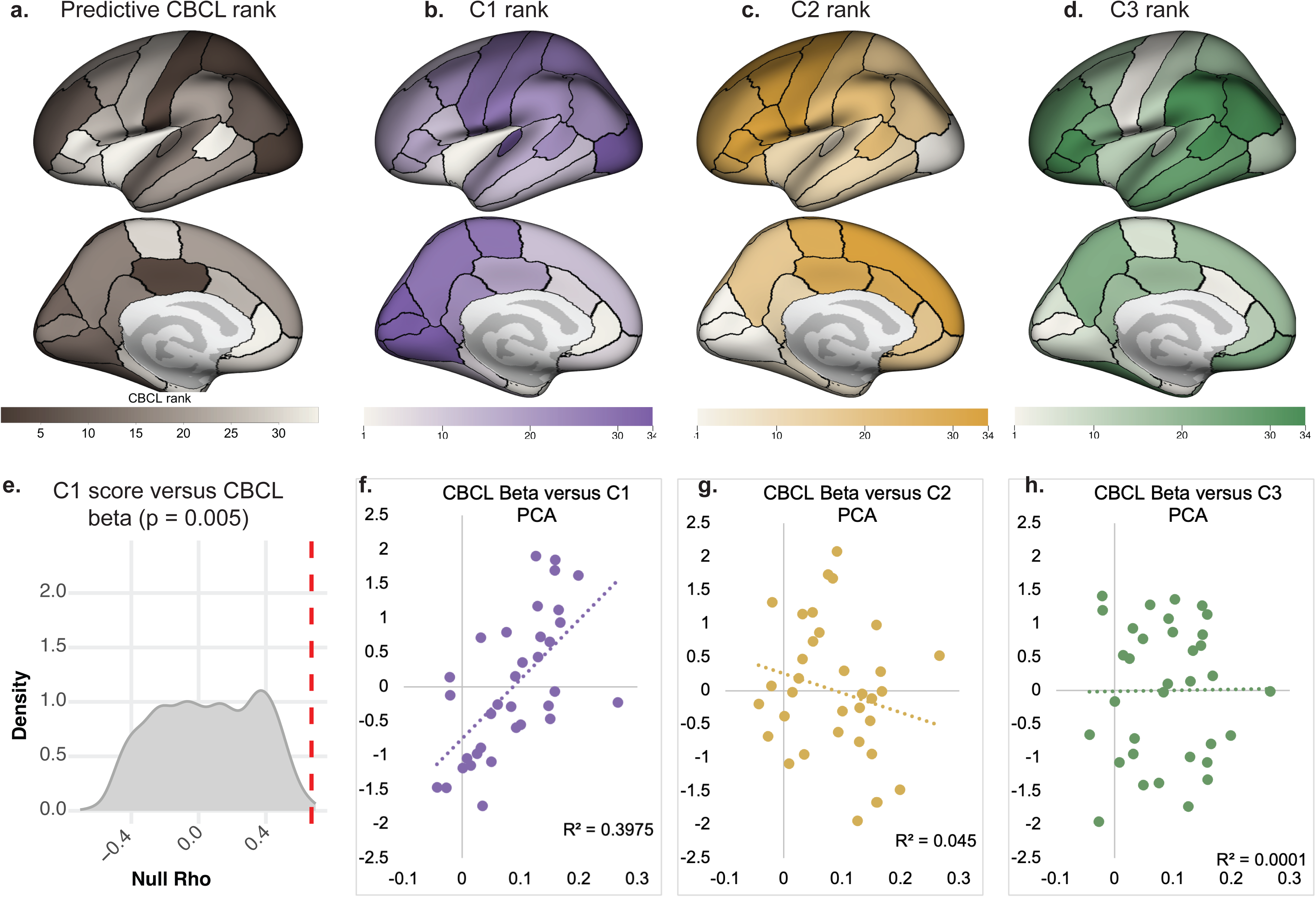
Convergence of predictive regions with spatiotemporal gene expression programs. Rank order of cortical regions based on CBCL total predictive beta **(a)** overlapped with rank order of regional enrichment for C1 (primarily prenatal) gene expression **(b)**, but not with rank order for C2 (primarily childhood, **c**) or C3 (primarily adolescent) enrichment **(d)**. **(e)** Spin testing indicated that rank order of C1 gene expression overlapped with that of CBCL predictive beta weights (p = 0.005). Using continuous data, regional prediction of CBCL total significantly correlated with regional enrichment for C1 expression **(f)**, but not of C2 **(g)** or C3 **(h)** gene expression. Right hemisphere data are not shown as C1, C2, and C3 expression profiles were only available for the left hemisphere.

## Discussion

These findings demonstrate patterned heterogeneity in maturation across the cortical surface that tracks with functional specialization, and predicts emergent psychopathology in adolescence. Convergent previous studies have established functional gradients across the cortical surface, including the SA axis^1^, that are generally consistent with the notion that more specialized (and evolutionarily newer) regions mature later in adolescence. Enabled by population-level neuroimaging and repeated measures across six years of study^16^, our work more precisely defines cortical maturation in terms of a deceleration in age-related cortical thinning, measurable by chronological differences in “break points” across brain regions. Break points occurred sequentially along the SA axis, providing a temporal framework for the hierarchical maturation of the cortex. Consistent with a previous study involving a substantially smaller sample but more frequent longitudinal imaging^23^, our findings support a sigmoid model of cortical maturation, characterized by an initial acceleration, followed by subsequent deceleration. These patterns were evaluated prospectively, by contrasting thinning contours apparent at ages 13-14 versus at ages 15-16. For example, while the initial acceleration was not observed in primary sensory regions likely because it occurred prior to age nine, age-related decelerations were seen across most of the surface by age 16. In contrast, select regions enriched for association cortex demonstrated only linear thinning over ages 9-16, suggesting that final maturation did not occur during this age range. These patterns were robust to variance in image quality, a key concern in pediatric neuroimaging studies; and were similar in male and female participants, with the latter showing steeper but temporally congruent declines in cortical thickness.

These findings help reconcile competing models for the relationship between psychopathology and cortical development. Many investigators have theorized that abnormal development of higher order association cortices must underlie the complex and varied symptoms which exist in psychiatric disease^24^. However, others have postulated that psychiatric symptoms arise from abnormalities in sensory regions, and these effects translate to symptoms as compromised sensory data moves through higher-order cortex^25^. We show that brain regions where increased thickness predicts mid-adolescent psychopathology overlap with those showing age-related decelerations in early adolescence, a process involving sensorimotor regions first, and association cortex later. This suggests that the location and function of the brain region where cortical development is altered may be less important than the timing of maturation in that region.

By extension, these findings raise the possibility of a relationship between eventual cortical thinning decelerations in long maturing regions and emergence of additional symptoms later in adolescence. Transdiagnostic staging models of psychiatric illness^14^ posit that early symptom manifestations, including depression, anxiety, or psychotic-like experiences, are often nonspecific and frequently comorbid. As illness progresses, such symptoms confer a broader vulnerability to increasingly severe psychopathology, which can be similar or dissimilar to symptoms encountered earlier. Consistent with this model, we found that early alterations in brain maturation predicted transdiagnostic psychiatric risk. Together, these findings suggest that earlier deviations in neurodevelopment may initially confer broad, rather than disorder-specific, psychiatric vulnerability, with greater phenotypic specificity emerging later in development. Further, the direction of this effect in early adolescence – that is to say, increased thickness predicting psychopathology risk – may shift later in development as additional regions mature. Indeed, longitudinal studies of cortical maturation in autism, which is characterized by a range of developmental psychopathology, demonstrate increased thickness before, and decreased thickness after adolescence compared to neurotypical controls^26^.

The enrichment of predictive regions for a gene set expressed primarily before birth implicates yet another asynchronous mechanism and reinforces the importance of longitudinal studies to decipher the biology of mental illness. Genes that have been associated with psychopathology risk in genome wide association studies are preferentially expressed in fetal life^27,28^, suggesting that they scaffold such risk for years before symptoms emerge. Similarly, cortical regions that show relatively early maturation in the present study are enriched for a network of prenatally-expressed genes, which in turn associate with increased risk for neurodevelopmental disorders^22^. Resolving specific mechanisms through which prenatally expressed genes ultimately influence cortical maturation and risk for emergent psychopathology in adolescence may provide critical insights into optimal developmental windows for early intervention and prevention, complementing work from ongoing longitudinal studies of brain development in the first decade of life^29^.

## Methods

### ABCD Study® and participants

This study used data from the ABCD Study® (Release 6.0), which enrolled 11,880 participants, mostly aged 9-10 at baseline, across 21 U.S. Institutional Review Board (IRB) approval for the ABCD Study is described in Auchter et al^30^. All parents provided written informed consent, and all youth gave informed assent. The present analyses used structural magnetic resonance imaging (sMRI) and behavioral data from baseline and the two, four, and six-year follow-up assessments (age range = 8.3 to 17.7); however, only the first half of the six-year follow-up data was available at the time of analysis.

### MRI acquisition

The sMRI data were acquired across study sites, using harmonized acquisition protocols across Siemens, Philips or GE 3T scanners. Details on image acquisition have been described in Casey et al.^16^. All T1-weighted images were acquired with a 256 × 256 matrix, slices = 176–225, repetition time (TR) = 6.31–2,500, echo time (TE) = 2–2.9, and 1 × 1 × 1-mm resolution. Minimally processed T1 volumes were available from the NIH Brain Development Cohorts Data Hub (https://www.nbdc-datahub.org/) for all but 44 participants. Participants were excluded if their sMRI scans failed cortical reconstruction.

### Image processing and cortical thickness

T1-weighted images were available for 11,752 participants at baseline, 8,093 at the two-year follow-up, 6,348 at the four-year follow-up, and 4,089 at the six-year follow-up. Image processing was performed as in Elyounssi et al^3^, including the following steps: 1) whole brain volumes were processed using FreeSurfer (version 7.1, http://surfer.nmr.mgh.harvard.edu/); 2) cortical thickness and total estimated intracranial volume were extracted from the Desikan-Killiany atlas (68 regions of interest (ROI), and whole left and right hemisphere); 3) SHN was calculated as an automated metric of quality control from the FreeSurfer aseg tabulated data output^3^. Based on Elyounssi et al.^3^, all analyses excluded scans with SHN ≥ 63. After quality control, 11,720 participants had at least one high-quality T1-weighted scan (SHN < 63), including 11,355 baseline, 7,895 two-year, 6,205 four-year, and 4,027 six-year scans (total number of scans: 29,482; see **Table S1**).

### Psychopathology

The parent-report Child Behavior Checklist (CBCL) was used as a measure of dimensional psychopathology, including three composite scores for total, internalizing, and externalizing symptoms. Age- and sex-corrected T-scores were used. Higher scores indicate more severe mental health problems, with T-scores ≥ 60 for composite scores representing clinically significant psychopathology^30^.

### Trajectories of age-related cortical thinning

We characterized cortical thickness trajectories in adolescents aged 8.3-17.7 years using longitudinal T1-weighted MRI data acquired across four time points. We used a tiered analysis approach, recognizing the importance of image quality control and trade-offs of cross-sectional versus longitudinal image acquisition^5^ in pediatric MRI scans. Head motion and other artifacts can systematically bias youth MRI studies^3^. However, this bias can be addressed through rigorous quality control. Following on our previous work,^3,6^ we used surface hole number (SHN), a validated automated metric that reliably excludes poor-quality images in the ABCD Study^3^. SHN can be used both to stratify MRI images based on thresholds derived from manually quality controlled images (e.g., SHN < 30 indicates high quality, SHN > 62 indicates poor quality) and as a scalar covariate in imaging analysis^3^. Further, longitudinal data better enables control for between-subject variation in unmeasured, time-invariant confounders that contribute to MRI measurements^7^.

As such, for analyses of age-related cortical thinning, we divided participants into (1) an optimized longitudinal dataset, comprising participants with high-quality scans (SHN < 30) available at all four time points, and complete covariates on age, sex, scanner, site, and family ID (*n* = 1,476); and (2) a non-overlapping, adequate remainder dataset included all remaining participants with at least one scan meeting the quality threshold (SHN < 63) and complete covariate data (*n* = 10,244). The optimized dataset was used for primary analyses of age-thinning trajectories, while the remainder dataset was used to assess reliability in a larger but lower-quality sample.

Linear mixed-effects models (LMMs; *lmerTest*) were implemented to examine cortical thickness trajectories across 68 regions defined by the Desikan–Killiany atlas. LMMs were chosen over other modeling approaches (such as generalized additive models) to facilitate categorization of thinning contours and to account for random effects (such as participant and family ID). Leveraging the serial data releases of ABCD, we initially examined thickness contours for the first three time points (baseline / most participants aged 9-10, year 2 / age 11-12, year 4 / age 13-14). This approach enabled us to distinguish linear and quadratic age-related thinning over this six-year interval, using two nested developmental models: a linear model including age, and a quadratic model including age and age^2^. Both models were adjusted for standard covariates of sex, intracranial volume, and SHN as fixed effects, and subject, family, scanner, and site as random effects to account for individual, family, scanner, and site-level variability. A preliminary quadratic designation (positive or negative) was assigned if the age^2^ term significantly predicted thickness after FDR correction across 68 regions (P_FDR_ < 0.05) and the Bayesian information criterion (BIC) was lower for the quadratic model; otherwise, a preliminary linear designation was assigned.

Next, we repeated the analysis including data from the fourth time point (year six / most participants aged 15-16). In this analysis, we could test for cubic patterns of thinning by adding a third nested model that also included age^3^. Again, the optimal developmental contour was determined based on the highest order statistically significant (P_FDR_ < 0.05) age term with the lowest BIC; otherwise, the next lower-order model was considered.

Finally, trajectory classifications were determined based on both the three- and four-time-point models. Four trajectory patterns were considered: early maturing (positive quadratic in both analyses), mid maturing (linear in the three-time-point and positive quadratic in the four-time-point model), fast maturing (cubic in the four-time-point model), and slow maturing (linear across all analyses).

To assess the robustness and generalizability of the findings in the optimized longitudinal dataset, we implemented the same analytical framework in the adequate remainder dataset. Consistency of cortical thickness trajectory classifications across the 68 regions between the two dataset was evaluated. Sensitivity analyses were conducted by adding fixed covariates for puberty stage (mean of available parent- and youth-reported scores), and income-to-needs ratio (INR, a global measure of socioeconomic status). Sex differences in cortical developmental trajectories were assessed by incorporating sex-by-age, -age^2^, and -age^3^ interaction terms into the best-fitting trajectory models as appropriate, again accounting for puberty stage and INR. FDR correction for multiple comparisons was implemented based on the number of regions with applicable age, age^2^, or age^3^ interaction terms.

### Breakpoint estimation in cortical thinning trajectories

To quantify the developmental timing of decelerations in cortical thinning trajectories, region-specific breakpoints were estimated as the age at which cortical thinning begins to slow. Breakpoints were identified using the R package *segmented*, which iteratively fits piecewise linear regression models, and estimates breakpoints that significantly improve model fit via maximum likelihood estimation. To improve estimation stability, a grid-search procedure was first implemented to determine optimal starting value. Candidate starting values ranged from age 9 (11 for fast maturing regions) to 17 years in 0.5-year increments. For each candidate, models were iteratively fitted until the change in residual sum of squares (RSS) was less than 1 × 10^-6^ or a maximum of 100 iterations was reached. The candidate yielding the lowest BIC was selected as the optimal starting value. Segmented regression was then refitted using the optimal starting value to obtain final breakpoint estimates for each cortical region using 1,000 bootstrap resamples, until the change in RSS was less than 1 × 10^-6^ or a maximum of 500 iterations was reached. Finally, for each region, breakpoint estimates with 95% confidence intervals (CIs), pre- and post-breakpoint slopes, cortical thickness at the breakpoint, and BIC were reported.

Breakpoint analyses were restricted to cortical regions with nonlinear developmental trajectories in the optimized longitudinal dataset. For early and mid maturing trajectories, all four time points were included in breakpoint estimation. For fast maturing regions, where baseline timepoints already showed early nonlinear development corresponding to accelerated cortical thinning, only the three follow-up assessments (two, four, and six-year) were used to estimate the second breakpoint, indicating deceleration.

### Spatial alignment of cortical thinning and sensorimotor–association axis

To test whether the timing of cortical developmental transitions aligns with the sensorimotor–association (SA) axis^31^, we examined the spatial correspondence between regional cortical thinning breakpoints and the log-transformed continuous hierarchical ordering of cortical regions along the SA axis. SA axis rank assignments were extracted from the framework derived by Sydnor et al.^31^. Statistical significance was assessed using spin testing to account for spatial autocorrelation across cortical regions^32^. Following the framework described by Alexander-Bloch et al.^32^, null models were generated by applying 5,000 spatial rotations to cortical maps on spherical representations of the cortex. Two-sided spin test p-values were calculated as the proportion of permuted correlations greater than or equal to the absolute observed correlation between breakpoint age and SA axis rank. Rotated surfaces across 68 DK^17^ cortical regions were generated separately within the left and right hemispheres.

### Relationships between cortical thickness and psychopathology

To examine the relationship between cortical thickness and psychopathology, we selected a sample of participants with adequate quality scans (SHN < 63), parent-report CBCL, and complete covariate data across timepoints baseline, year two, and year four. Data from year six was not included due to significantly smaller sample size.

The longitudinal relationship between cortical thickness and dimensional psychopathology was assessed using three competing temporal models, using thickness and CBCL total data from baseline, year two, and year four. A *concurrent* model (n = 10,402) tested same-visit associations between cortical thickness and CBCL scores at each time point. A *predictive* model (n = 9,474) tested whether early cortical thickness was associated with later CBCL, including baseline cortical thickness predicting two-year and four-year CBCL scores and two-year cortical thickness predicting four-year CBCL scores, respectively. A *delayed* model (n = 8,481) tested the reverse temporal direction, assessing whether earlier CBCL symptoms were associated with later cortical thickness. Models were first tested on the mean cortical thickness using linear mixed-effects models (LMMs; *lmerTest*, R v4.4.0) with the Wald test to evaluate effects across timepoint pairs, while accounting for subject-level information. The Wald test was chosen because it accounts for simultaneous use of repeated data (e.g., baseline thickness as predictors of both year 2 and year 4 thickness). All models included fixed effects of age, sex, SHN, ICV, INR, puberty stage, and random effects of family ID, scanner, and site. Only the predictive model was statistically significant at the mean cortical thickness value after FDR correction across three tests (P_FDR_ < 0.05), and this model was carried forward to subsequent analyses.

Predictive relationships between psychopathology and cortical thinning were then tested across the 68 DK atlas ROIs using the joint Wald test, controlling for fixed effects of age, sex, SHN, ICV, INR, puberty stage, and random effects of family ID, scanner, and site. For thoroughness, we reported regions which had a statistically significant predictive relationship after FDR correction across 68 tests (P_FDR_ < 0.05) and those which were nominally significant (P < 0.05). Next, we examined specific year pairs of predictive relationships, including baseline-year two, baseline-year four, and year two-year four, based on both FDR correction across 68 tests (P_FDR_ < 0.05) and nominal significance (P < 0.05).

Spin testing was used to assess whether regions which showed a predictive relationship with CBCL were enriched for linear versus non-linear trajectories, beyond what would be expected by chance under spatial autocorrelation. Null models were generated by applying 5,000 spatial rotations to cortical maps on spherical representations of the cortex. Two-sided spin test p-values were calculated as the proportion of permuted absolute differences greater than or equal to the absolute observed difference. Rotated surfaces across 68 DK cortical regions were generated separately within the left and right hemispheres.

### Case-control analysis

Post hoc analyses were performed to test whether cortical thickness in regions showing significant predictive associations differed between children with low versus clinically significant CBCL groups at the corresponding outcome time point. Analyses were conducted separately for each prediction interval: baseline cortical thickness predicting 4-year CBCL scores, 2-year cortical thickness predicting 4-year CBCL scores, and baseline cortical thickness predicting 2-year CBCL scores. For each interval, participants were ranked by their CBCL T-score and divided into tertiles; the lowest tertile defined the low-CBCL group, whereas clinically significant psychopathology was defined as a CBCL total T-score ≥60^33^. To improve group comparability, propensity-score matching (R package *MatchIt*^34^) was performed based on age, sex, ICV, SHN, INR, pubertal stage, site, and scanner at the cortical imaging time point. Participants were matched using 1:1 nearest-neighbor matching without replacement, with the clinically significant group as the target population and a standardized caliper of 0.2^35^. Covariate balance before and after matching was quantified using standardized mean differences, with an absolute standardized mean difference <0.1 used as an indicator of adequate balance.

For each prediction interval, conditional logistic regression was fitted to examine association from cortical thickness to the odds of clinically significant CBCL group compared to low-CBCL group at the corresponding outcome time point. Models were stratified by matched pairs to preserve the matched design, with variance estimates clustered at the family level to account for within-family correlation. Mean cortical thickness for each group, odds ratios (ORs) with 95% confidence intervals (CIs), and corresponding P_FDR_ values were reported.

### Genetic patterning

C1, C2, and C3 genes sets were previously derived using diffusion map embedding of Allan Human Brain Atlas (AHBA) data^36^ per^22^. Loading for expression of C1, C2, and C3 genes was calculated for each of the 34 left-sided cortical regions in the DK atlas, as previously determined by^22^. Spin tests (as above) determined the degree of spatial overlap between C1, C2, and C3 loading and predictive betas (for the overall relationship between thickness and future psychopathology) for each left-sided cortical region. Only left-sided expression data were used due to limited availability of right hemisphere data in the AHBA.

## Data availability

Data used in this study were obtained from the Adolescent Brain Cognitive Development (ABCD) Study, release 6.0, through the National Brain Development Cohort Data Hub (https://www.nbdc-datahub.org/). Participant-level data are available to qualified researchers through the NDA’s controlled-access application process and subject to its data use requirements.

## Code availability

Code used to perform the analyses is available to editors and reviewers upon request during peer review and will be publicly available at [REPOSITORY] upon publication.

## Extended Data Legends

**Extended Data Fig. 1:**
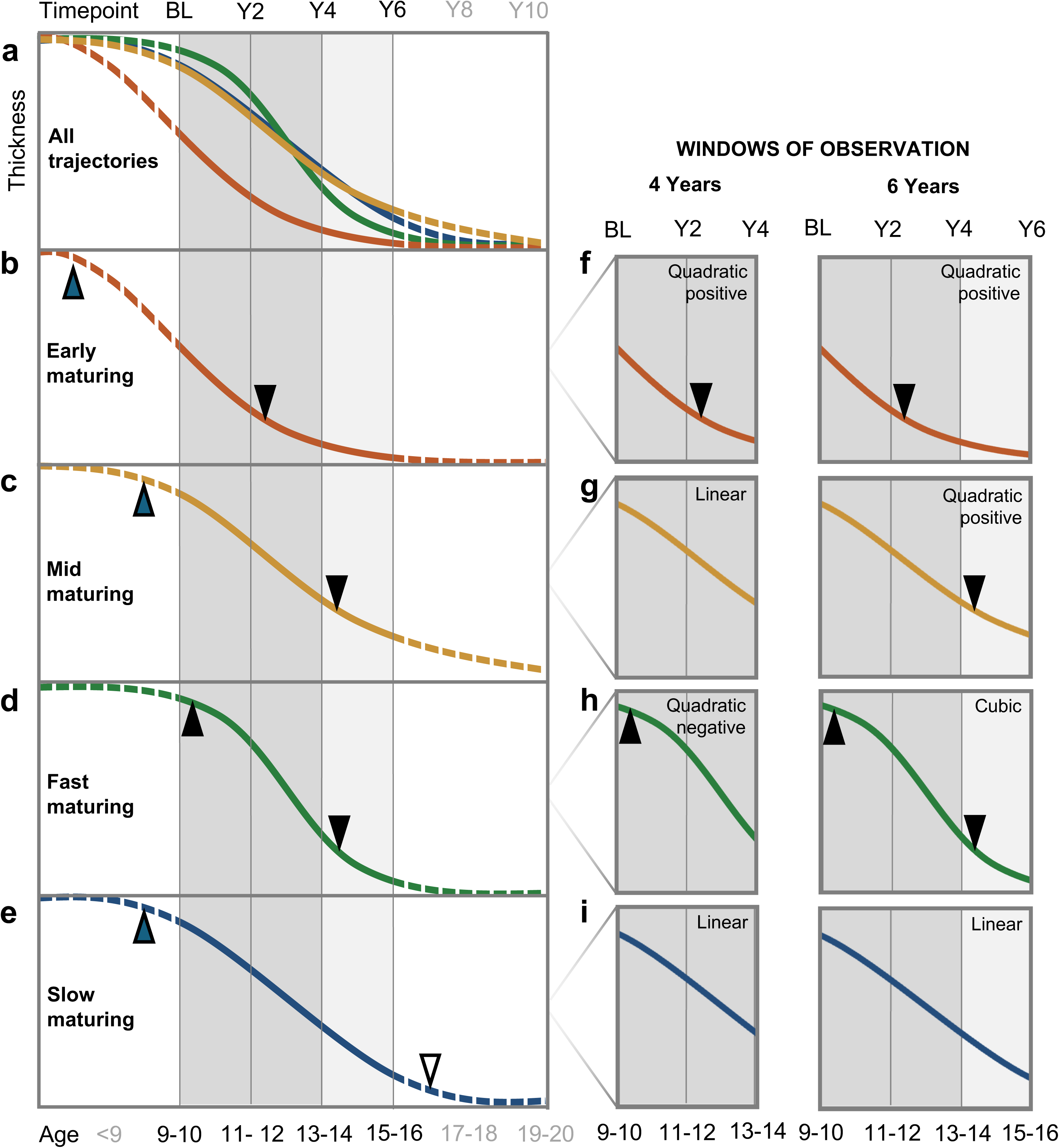
A sigmoid model of age-related cortical thinning, with regionally specific onsets and durations, accounts for all observed data (solid curves, in **(a)** through **(e)**). Upward triangles indicate acceleration phases, and downward triangles indicate deceleration phases. While fast maturing regions **(d)** demonstrate the full sigmoid contour, early **(b)**, mid **(c)**, and slow **(e)** maturing regions would have shown initial accelerations before the present observation window of ages 9 to 16 (open triangles). Further, slow maturing regions would not show thickness decelerations until after age 16 (open triangles); these decelerations may be detected at future ABCD timepoints. Consistent with observed data, apparent contours can depend on the timing and duration of the observation window (**f** through **i**) as well as image quality (see also Supplemental Figure 1).

**Extended Data Fig. 2:**
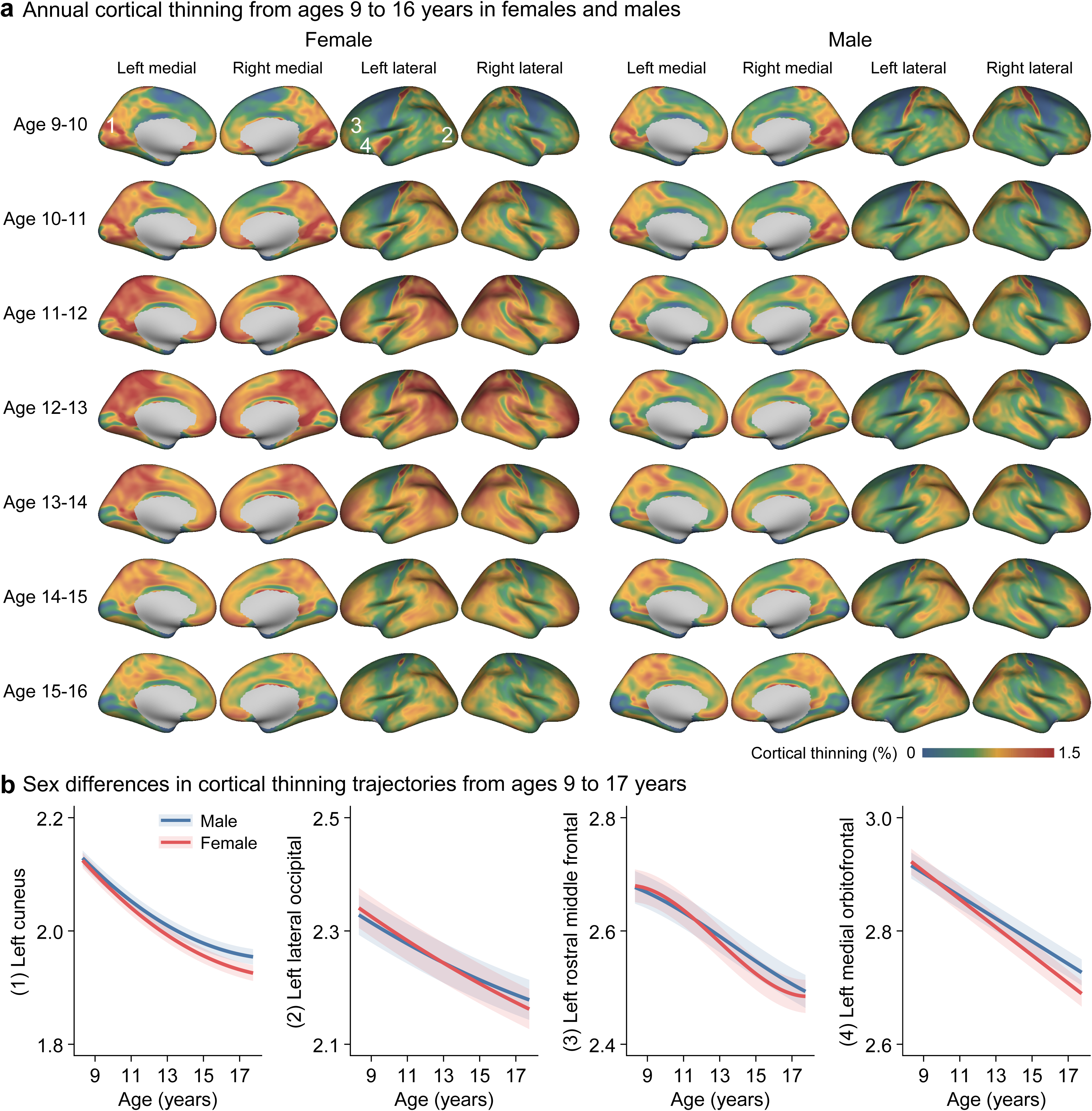
(**a**) Vertex-wise maps of annual cortical thinning across ages 9–16 years, after correction for standard covariates other than sex assigned at birth (SAB), separated by SAB. See Fig. 1 for comparison. Maps show annual cortical thinning percentages for each age interval from 9–10 to 15–16 years. Warmer colors indicate more annual cortical thinning, whereas cooler colors indicate less thinning. (**b)** SAB-stratified age-related cortical thinning trajectories are shown for regions that are representative of different thinning trajectories. Clouds indicate 95% CI.

**Extended Data Fig. 3:**
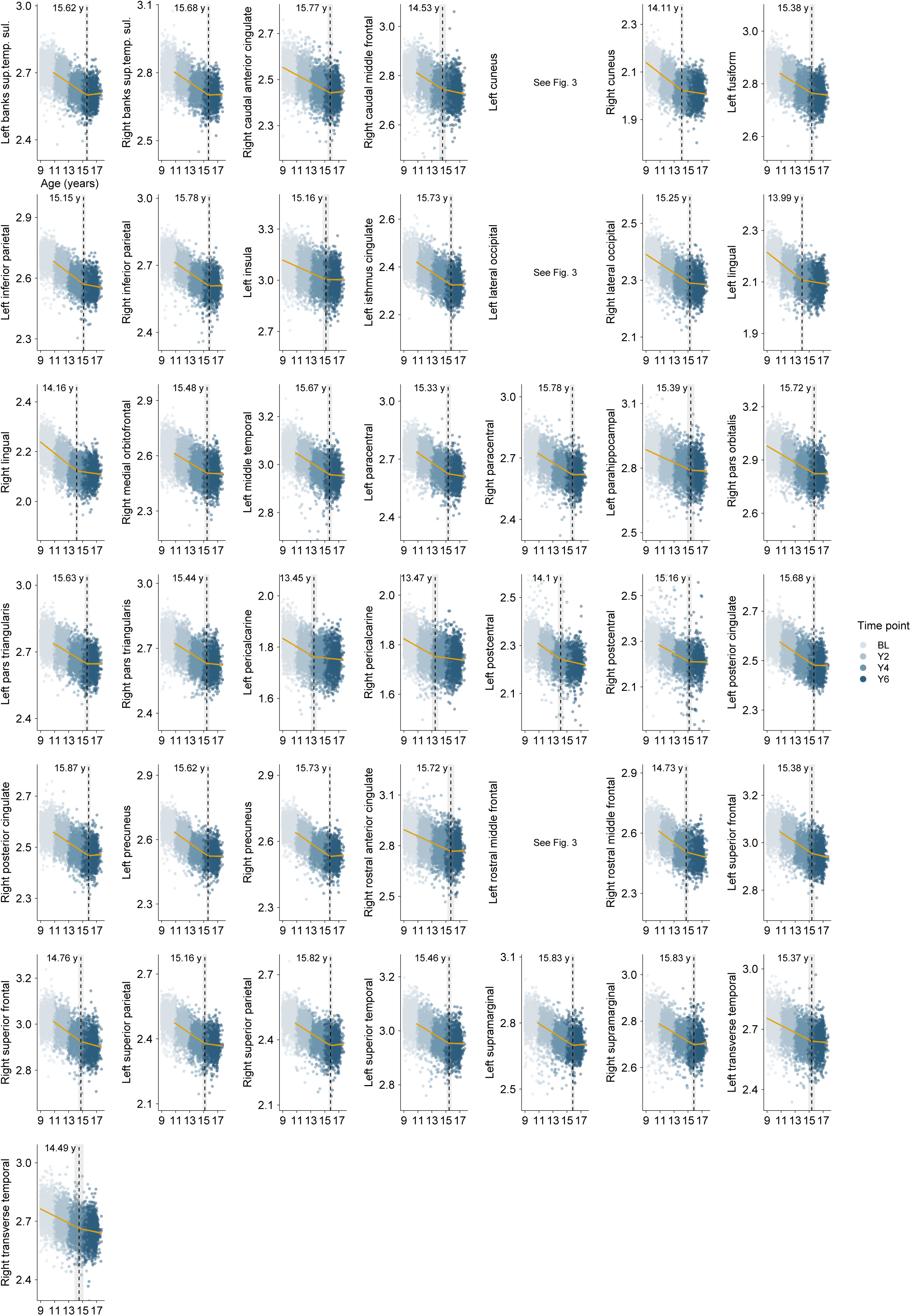
Breakpoint estimation for representative cortical thickness trajectories for 40 remaining non-linear regions (see also Fig. 3). Regions are ordered alphabetically. Grey vertical lines indicate estimated breakpoint age with 95% confidence intervals (CIs). Orange lines represent segmented linear model fits before and after the breakpoint. Each point represents an individual observation, color-coded by imaging wave.

**Extended Data Fig. 4:**
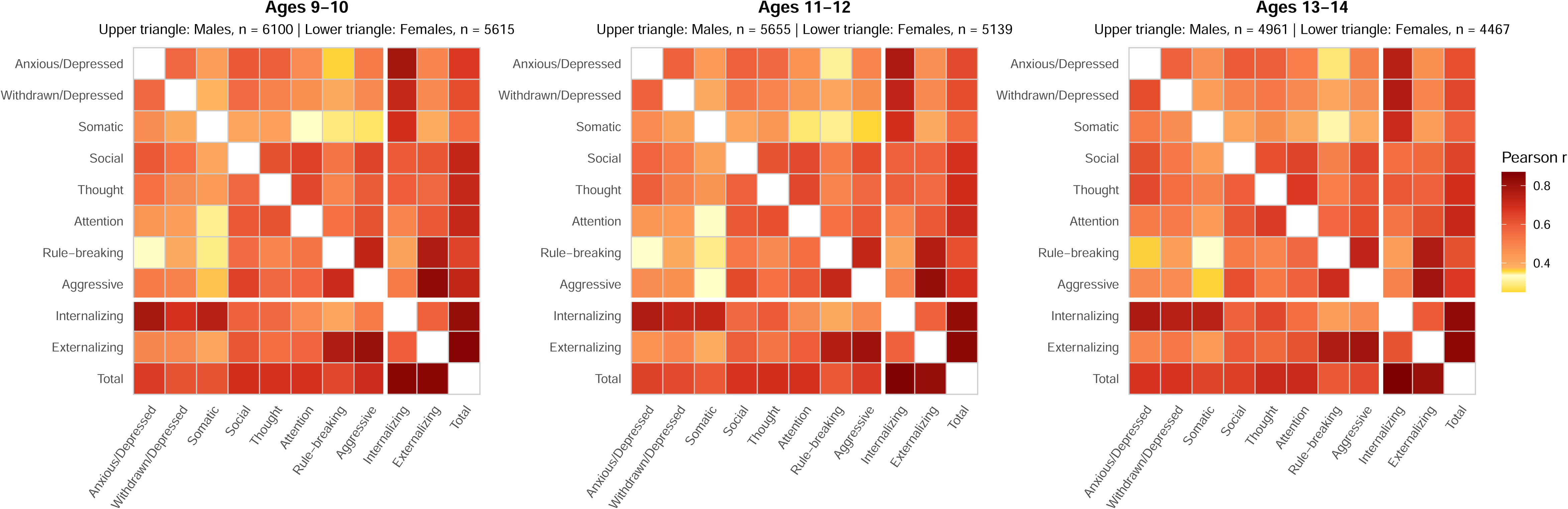
Cross-correlations of CBCL narrow- and broadband scores at baseline (ages 9-10), year 2 (ages 11-12), and year 4 (ages 13-14).

**Extended Data Fig. 5:**
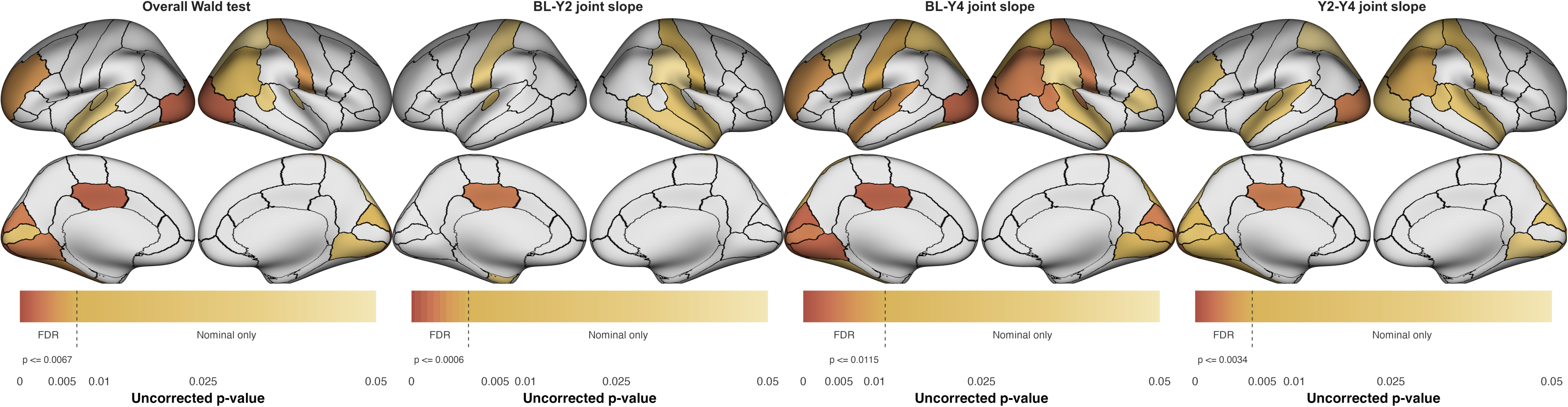
**a)** Omnibus results showing regions that predicted CBCL total score two years later (including baseline to year 2, baseline to year 4, and year 2 to year 4). **(b)** Regions where baseline thickness predicted CBCL total at year 2. **(c)** Regions where thickness at year 2 predicted CBCL total at year 4. See also Fig. 4b for regions where baseline thickness predicted CBCL total at year 4.

**Extended Data Fig. 6:**
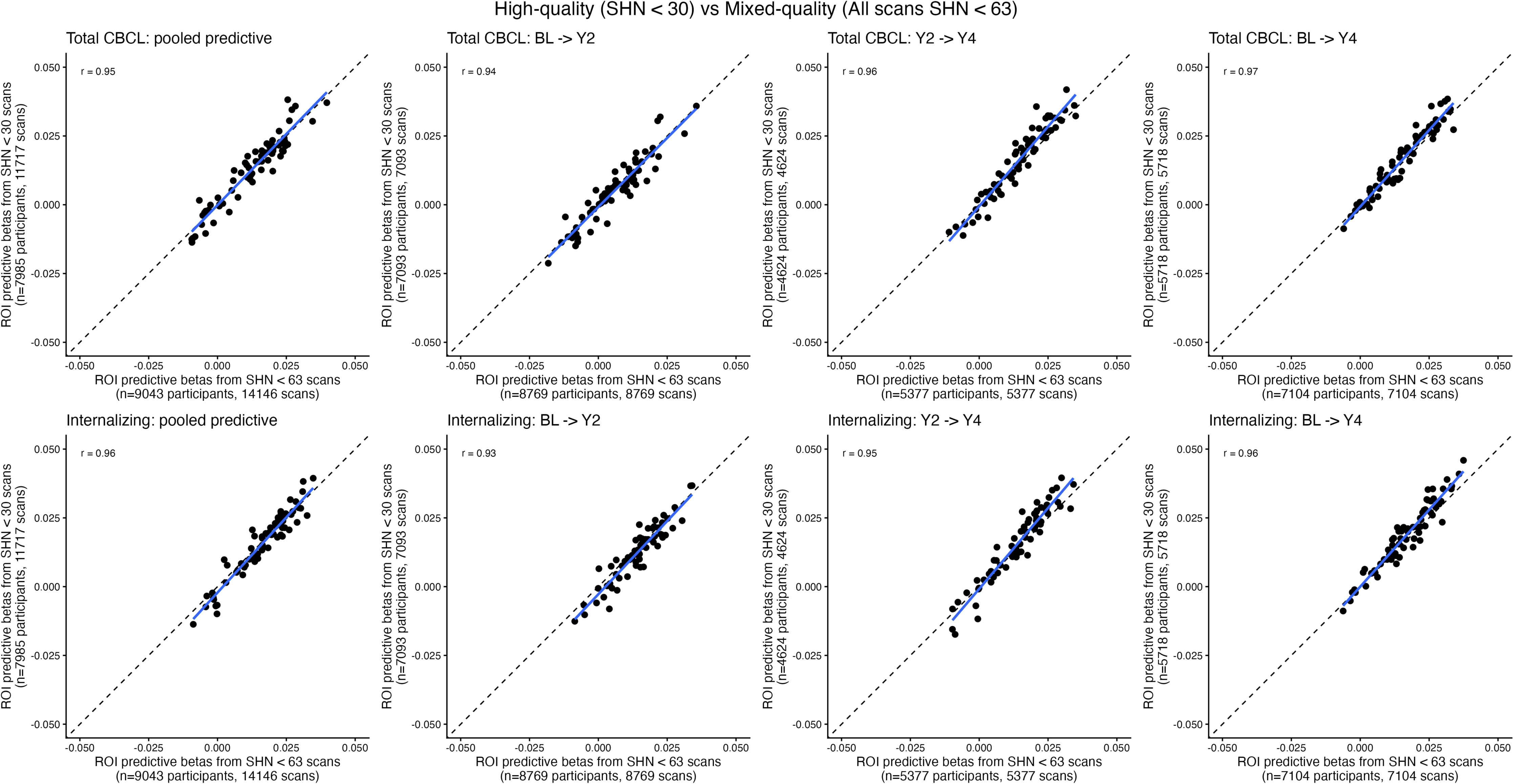
Reproducibility of thickness predicting psychopathology (beta) at different intervals, comparing entire available sample (x-axis) with subsample that have high-quality scans (SHN < 30, y-axis). Each data point is one of 68 regions of interest from the D-K atlas. Blue lines indicate best-fit, and dashed lines indicate unity. Top row represents CBCL total scores, and bottom row represents CBCL internalizing symptom scores.

## Author Contributions

**Conception:** Clauss, Zhi, O’Dor, Flook, Linneman, Kritzer, Holt, P. Lee, Dunn, Satterthwaite, Barch, Gilman, Vertes, Bullmore, Roffman

**Design:** Clauss, Zhi, Dear, Kunitoki, Hughes, Vinke, H. Lee, Doyle, Roffman

**Data analysis:** Clauss, Zhi, Perdomo, Kunitoki, Blum, Kane, Elyounssi, Cooper, Laurent, Bazer

**Drafting/revision:** All authors

**Approved final manuscript:** All authors

## Additional Information

Supplementary Information is available for this paper.

Reprints and permissions information are available at www.nature.com/reprints.

