## Supplementary figures and images for "Inflections in adolescent cortical maturation signal psychopathology risk"

### Supplemental Figure 1

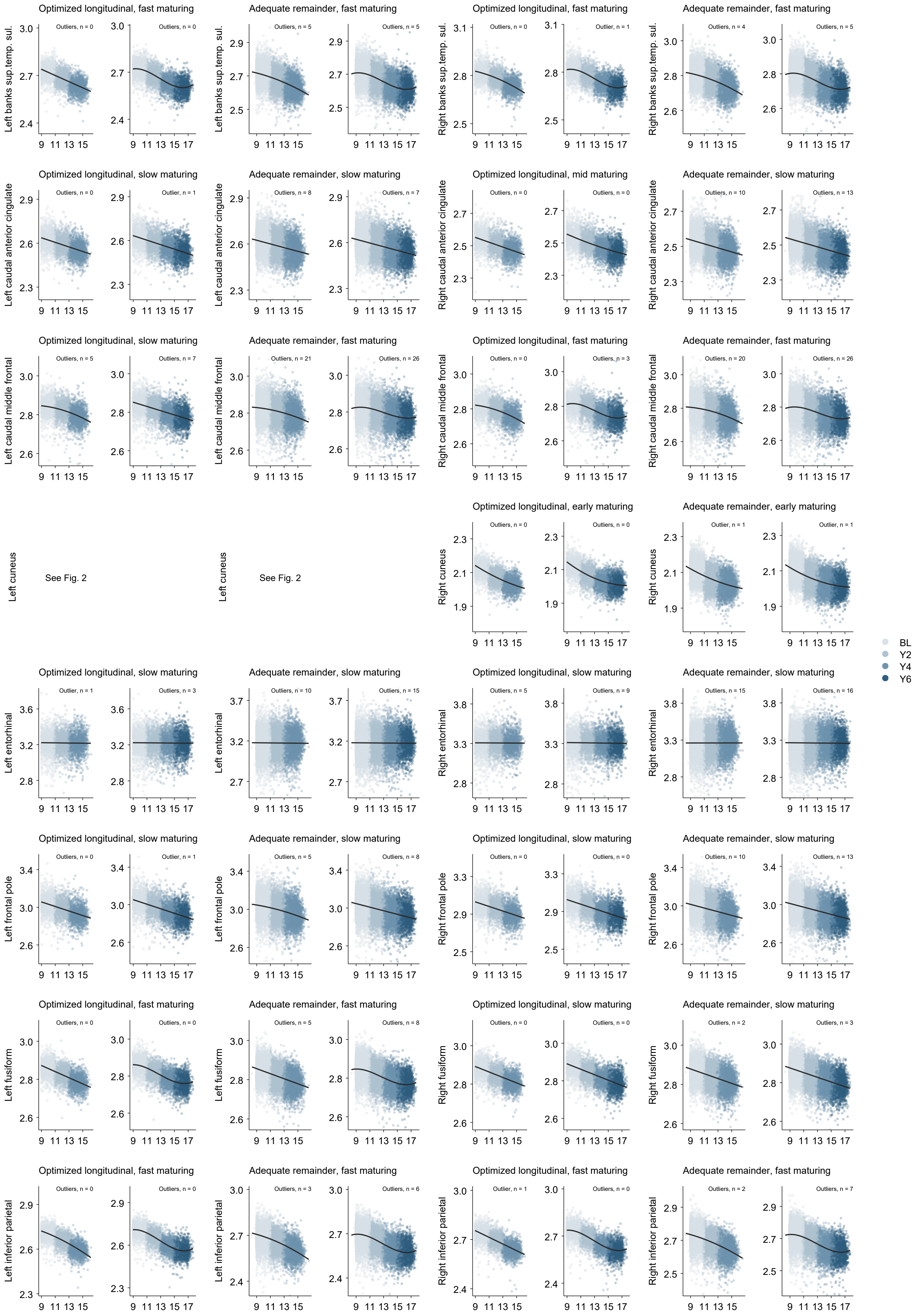

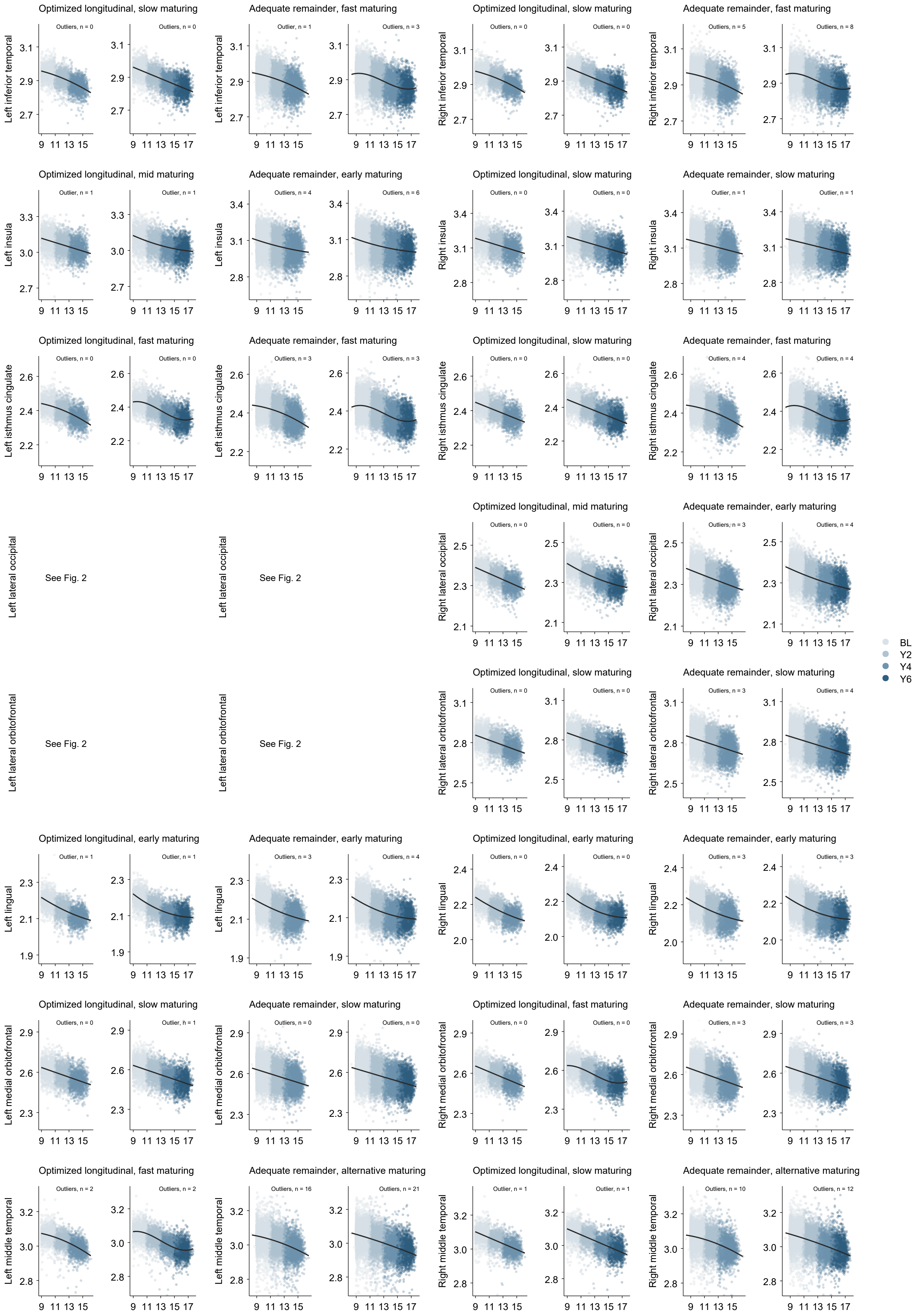

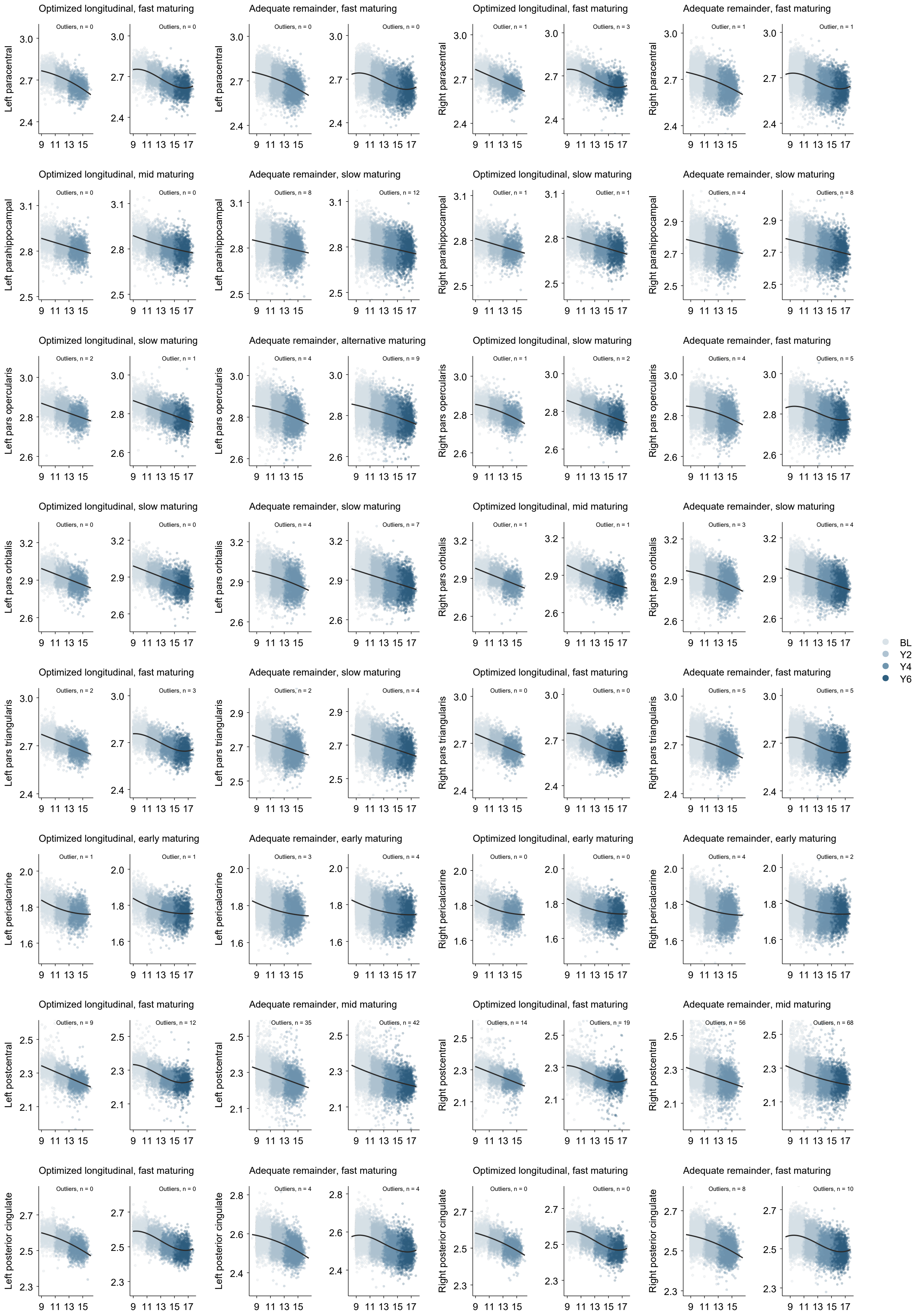

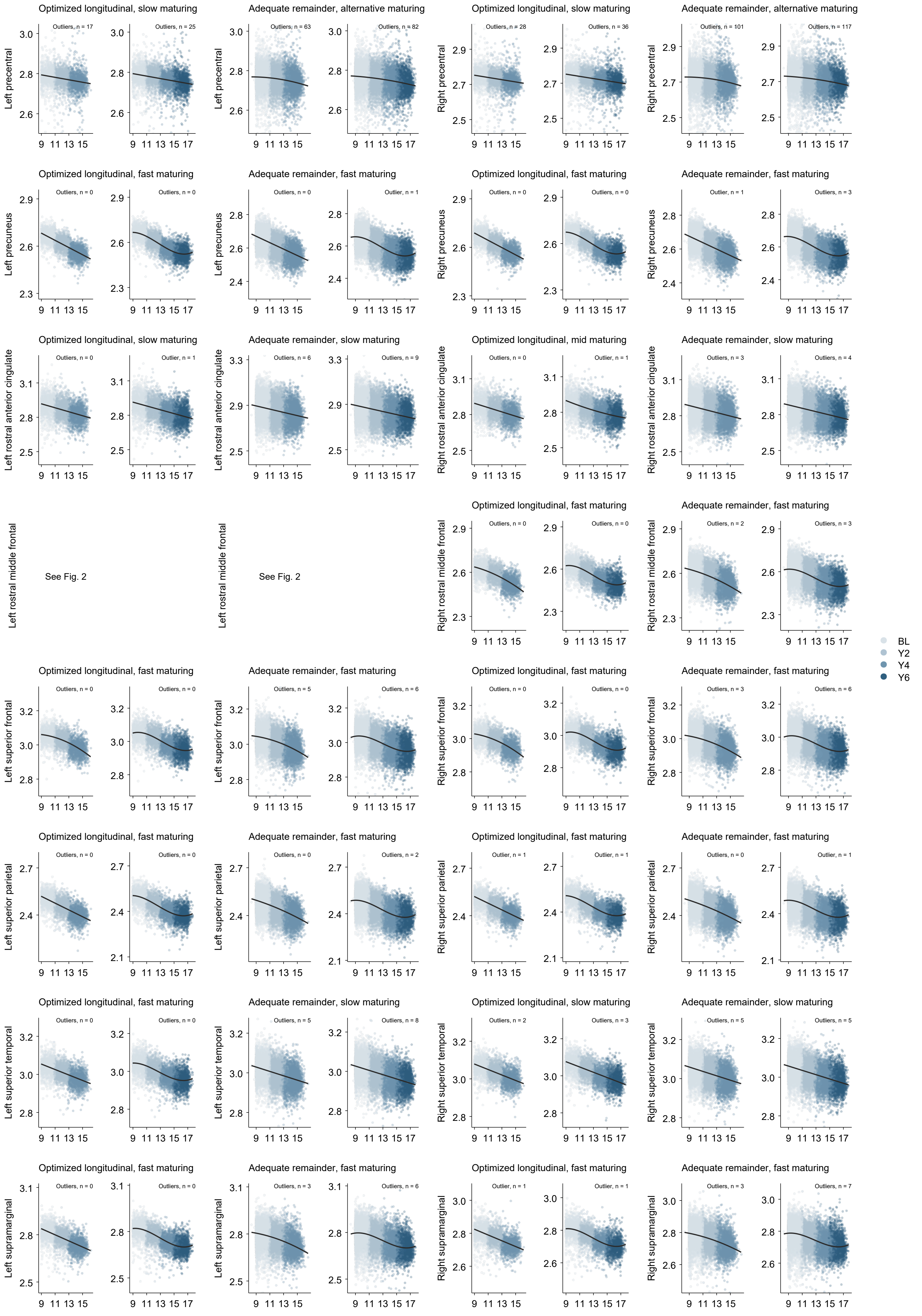

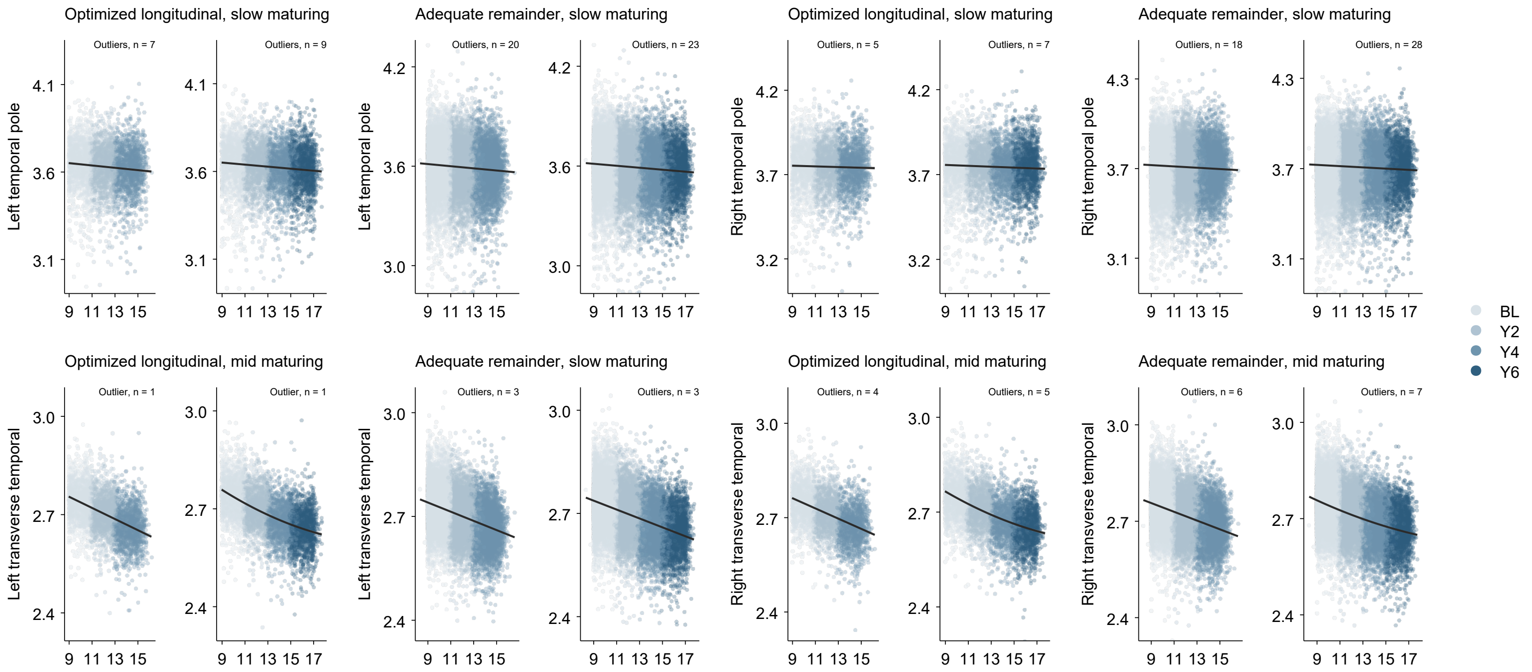
